# Genetic Determinants Underlying the Progressive Phenotype of β-lactam/β-lactamase Inhibitor Resistance in *Escherichia coli*

**DOI:** 10.1101/2023.05.24.542208

**Authors:** William C Shropshire, Hatim Amiji, Jordan Bremer, Selvalakshmi Selvaraj Anand, Benjamin Strope, Pranoti Sahasrabhojane, Marc Gohel, Samuel Aitken, Sarah Spitznogle, Xiaowei Zhan, Jiwoong Kim, David E Greenberg, Samuel A Shelburne

**Affiliations:** Department of Infectious Diseases, Infection Control, and Employee Health, The University of Texas MD Anderson Cancer Center, Houston, TX, USA; Frank H. Netter MD School of Medicine, Quinnipiac University, Hamden, CT, USA; Program in Diagnostic Genetics and Genomics, MD Anderson Cancer Center School of Health Professions, Houston, TX, USA; Division of Pharmacy, The University of Texas MD Anderson Cancer Center, Houston, TX, USA; Quantitative Biomedical Research Center, Peter O’Donnell Jr. School of Public Health, University of Texas Southwestern Medical Center, Dallas, TX, 75390; Department of Internal Medicine, UT Southwestern, Dallas, TX 75390, USA; Department of Microbiology, UT Southwestern, Dallas, TX 75390, USA; Department of Genomic Medicine, The University of Texas MD Anderson Cancer Center, Houston, TX, USA

## Abstract

Currently, whole genome sequencing (WGS) data has not shown strong concordance with *E. coli* susceptibility profiles to the commonly used β-lactam/β-lactamase inhibitor (BL/BLI) combinations: ampicillin-sulbactam (SAM), amoxicillin-clavulanate (AMC), and piperacillin-tazobactam (TZP). Progressive resistance to these BL/BLIs in absence of cephalosporin resistance, also known as extended-spectrum resistance to BL/BLI (ESRI), has been suggested to primarily result from increased copy numbers of *bla*_TEM_ variants, which is not routinely assessed in WGS data. We sought to determine whether addition of gene amplification could improve genotype-phenotype associations through WGS analysis of 147 *E. coli* bacteremia isolates with increasing categories of BL/BLI non-susceptibility ranging from ampicillin-susceptible to fully resistant to all three BL/BLIs. Consistent with a key role of *bla*_TEM_ in ESRI, 112/134 strains (84%) with at least ampicillin non-susceptibility encoded *bla*_TEM_. Evidence of *bla*_TEM_ amplification (i.e., *bla*_TEM_ gene copy number estimates > 2×) was present in 40/112 (36%) strains. There were positive correlations between *bla*_TEM_ copy numbers with minimum inhibitory concentrations (MICs) of AMC and TZP (*P*-value < 0.05), but not for SAM (*P*-value = 0.09). The diversity of β-lactam resistance mechanisms, including non-ceftriaxone hydrolyzing *bla*_CTX-M_ variants, *bla*_OXA-1_, as well as *ampC* and *bla*_TEM_ strong promoter mutations, were greater in AMC and TZP non-susceptible strains but rarely observed within SAM and AMP non-susceptible isolates. Our study indicates a comprehensive analysis of WGS data, including β-lactamase encoding gene amplification, can help categorize *E. coli* with AMC or TZP non-susceptibility but that discerning the transition from SAM susceptible to non-susceptible using genetic data requires further refinement.

**Importance:** The increased feasibility of whole genome sequencing has generated significant interest in using such molecular diagnostic approaches to characterize difficult-to-treat, antimicrobial resistant (AMR) infections. Nevertheless, there are current limitations in the accurate prediction of AMR phenotypes based on existing AMR gene database approaches, which primarily correlate a phenotype with the presence/absence of a single AMR gene. Our study utilized a large cohort of cephalosporin-susceptible *E. coli* bacteremia samples to determine how increasing dosage of narrow-spectrum β-lactamase encoding genes in conjunction with other diverse BL/BLI genetic determinants contribute to progressively more severe BL/BLI phenotypes. We were able to characterize the complexity of the genetic mechanisms underlying progressive BL/BLI resistance including the critical role of β-lactamase encoding gene amplification. For the diverse array of AMR phenotypes with complex mechanisms involving multiple genomic factors, our study provides an example of how composite risk scores may improve understanding of AMR genotype/phenotype correlations.

## Introduction

Rapid and accurate characterization of antimicrobial resistant (AMR) infections is necessary to combat their increasing threat to public health (1, 2).There has been recent interest both from a clinical and research standpoint in using whole genome sequencing (WGS) to predict antimicrobial susceptibility patterns as molecular diagnostic approaches become more feasible (3, 4). When AMR is driven by a single gene, such as *bla*_CTX-M_ for an extended-spectrum β-lactamase phenotype, AMR database query approaches generally detect a strong concordance between WGS predicted and observed phenotypes (5). However, discordant genotype-phenotype predictions can occur due to complicated resistance mechanisms that involve multiple contributing genetic factors (6). Such issues have been consistently observed when trying to use WGS to predict the antimicrobial susceptibility pattern of β-lactamase/β-lactamase inhibitor (BL/BLI) combinations (5, 7, 8).

Although recent years have seen the introduction into clinical practice of such novel BL/BLI combinations as ceftolozane/tazobactam and ceftazidime/avibactam, the vast majority of BL/BLIs currently used are ampicillin-sulbactam (SAM), amoxicillin-clavulanate (AMC), and piperacillin-tazobactam (TZP) (9–11). Given the clinical impact of *Escherichia coli* and its highly varied susceptibility profile to these three BL/BLI combinations, *E. coli* is among the most well studied organisms in terms of BL/BLI resistance (8, 12–21). Cephalosporinases and carbapenemases that hydrolyze broad spectrum β-lactams generally also inactivate SAM, AMC, and TZP (22). There has been increasing interest in characterizing cephalosporin-susceptible *E. coli* strains with varying susceptibility patterns amongst these three common BL/BLI combinations (13–16). Nevertheless, the characterization of these increasingly more severe BL/BLI resistant phenotypes using WGS to detect potential associations in well-defined cohorts are less understood.

To date, most studied *E. coli* ceftriaxone-susceptible, BL/BLI resistant strains harbor class A or class D beta-lactamases such as *bla*_TEM-1B_ or *bla*_OXA-1_ (13–15). It has been suggested that the BL/BLI resistance amongst such strains evolves in a gradient fashion in order from ampicillin-sulbactam non-susceptible (SAM-NS) to amoxicillin-clavulanate non-susceptible (AMC-NS) to piperacillin-tazobactam non-susceptible (TZP-NS) (13, 23). Based on studies in a limited number of strains under laboratory passage as well as in clinical isolates, *bla*_TEM_ amplification has been proposed as a major mechanism of what has been called extended-spectrum resistance to BL/BLIs or ESRI (13-15, 18, 21). However, there are many other genetic determinants that can contribute to reduced susceptibility to BL/BLIs such as augmented expression of *bla*_TEM_ due to promoter mutations (24, 25), de-repression of the chromosomal *ampC* gene (26, 27), decreased outer membrane permeability (20, 28), inhibitor resistant TEM variants (29), as well as CTX-M enzymes with decreased cephalosporin affinity but increased TZP hydrolysis activity (30). Furthermore, it is not clear how these BL/BLI genetic determinants contribute to BL/BLI resistance across the full ESRI spectrum.

There has been in-depth examination of *E. coli* BL/BLI resistance mechanisms with regards to each respective BL/BLI combination. For example, Noguchi et al. used a phenotypic approach and found that hyperproduction of TEM-1 and altered cell membrane permeability accounted for a large proportion of SAM resistance (20). Conversely, WGS based analyses of AMC and TZP resistance showed a complex variety of mechanisms including *bla*_TEM-1_ overexpression as well as increased copy numbers of various *bla* encoding enzymes (8, 16). A critical missing piece in understanding the role of genetic analyses in BL/BLI resistance is the lack of WGS investigation of a large number of cephalosporin-susceptible *E. coli* isolates with varying BL/BLI resistance profiles. Thus, we performed WGS of 147 ceftriaxone-susceptible *E. coli* which had BL/BLI resistance phenotypes ranging from ampicillin (AMP) susceptible to TZP non-susceptible. We sought to determine whether genetic analysis of known BL/BLI resistance mechanisms could be used to classify amongst the various phenotypic ESRI categories. Our data suggest that progressive ESRI is driven in part by *bla*_TEM_ amplification but that many complex mechanisms detectable by WGS analysis also contribute to AMC and TZP non-susceptibility. The lack of ability to separate SAM susceptible from SAM non-susceptible using WGS data, including analysis of *bla*_TEM_ copy number, indicates that improvements of these genotype/phenotype correlations may require alternative approaches to current AMR genetic database query methods.

## Results

### Distribution of ESRI groups amongst CRO-S *Escherichia coli* BSI isolates

We identified 1,026 *E. coli* bloodstream isolates from May 1^st^, 2015, to April 30^th^, 2020. Of these, 389 (38%) had a ceftriaxone (CRO) MIC ≥ 4 mg/L. Consistent with a high rate of BL/BLI resistance among CRO resistant isolates, 91% and 76% were non-susceptible to SAM and AMC respectively. We selected the remaining 637 (62%) *E. coli* isolates in our sampling frame with a CRO MIC < 4 mg/L to study cephalosporin susceptible, BL/BLI associated resistance mechanisms (Fig. S1A). We grouped isolates by their susceptibility patterns according to CLSI guidelines as follows: β-lactam pan-susceptible (PAN-S; Group 1), ampicillin non-susceptible (AMP-NS; Group 2), ampicillin-sulbactam non-susceptible (SAM-NS; Group 3), amoxicillin-clavulanate non-susceptible (AMC-NS; Group 4), and piperacillin-tazobactam non-susceptible (TZP-NS; Group 5) as shown in **Fig. 1**.

**FIG 1.**
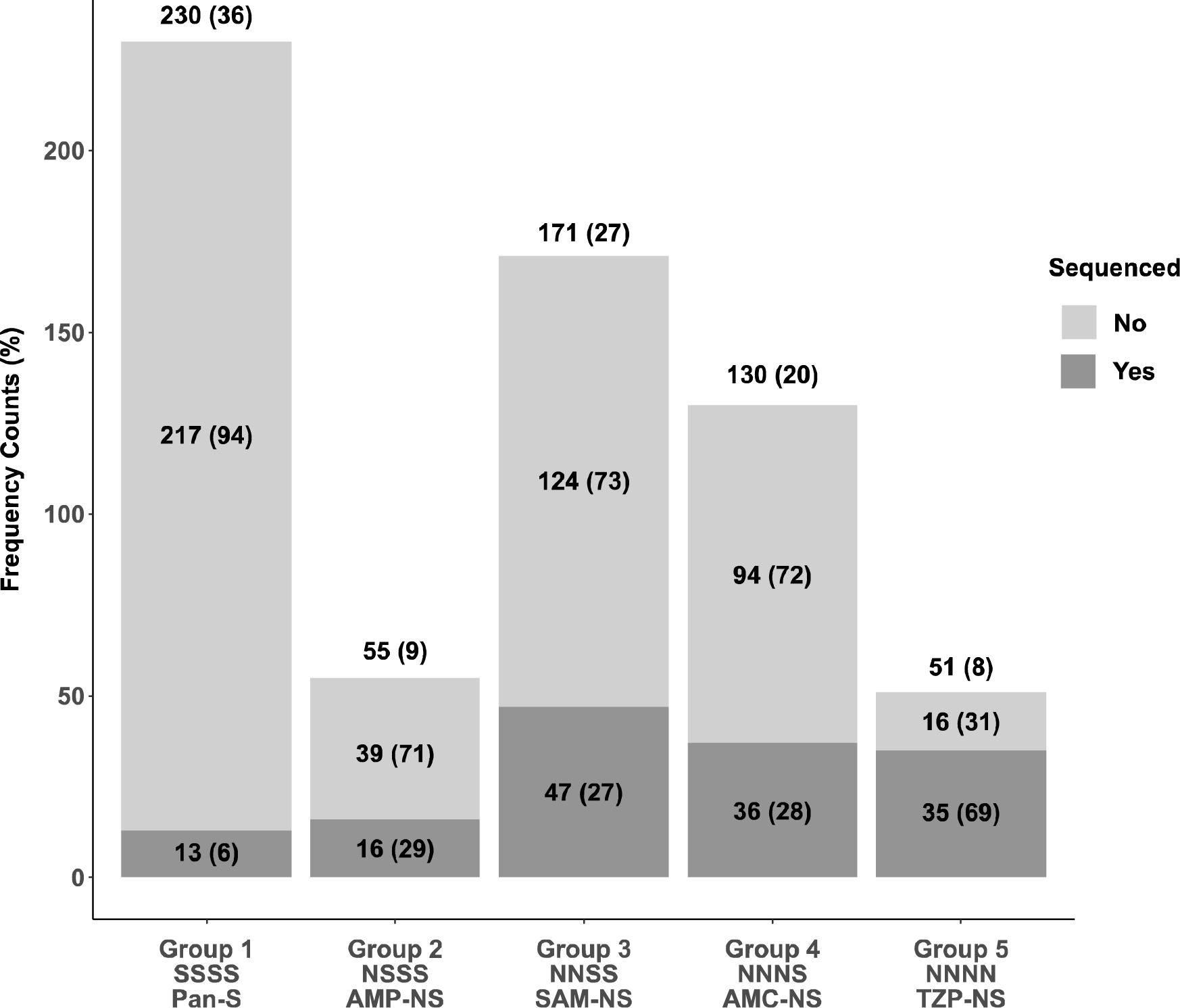
Summary of ceftriaxone susceptible (*i.e.*, CRO MIC < 4 mg/L) *Escherichia coli* bloodstream isolates (n=637) stratified by increasing BL/BLI resistant phenotype (Group 1 – 5) along with number of sequenced isolates from each group. Unique BSI samples are grouped into progressively more resistant BL/BLI phenotypes from pan-susceptible (Group 1) to resistant to all four drug combinations in our study (Group 5). Frequency counts above each bar give group totals with percentage of total population in parenthesis [*e.g.*, there were 230 Group 1 isolates out of the total 637 (36%)]. We further split groups into sequenced (n= 147) (dark gray) vs non-sequenced (light gray) with frequency counts and percentages of each respective group labelled [*e.g.*, 13 out of 230 group 1 isolates were sequenced (6 %)]. N = Non-susceptible; S = Susceptible; Pan-S = pan-susceptible; AMP-NS = ampicillin non-susceptible; SAM-NS = ampicillin-sulbactam non-susceptible; AMC = amoxicillin-clavulanate non-susceptible; TZP-NS = piperacillin-tazobactam non-susceptible.

The plural majority (230/637; 36%) of CRO susceptible *E. coli* BSI isolates were pan-susceptible to the studied β-lactams (Group 1). The next most common isolates were SAM-NS (Group 3; 27%) and AMC-NS (Group 4; 20%) whereas the lowest frequencies were found within AMP-NS (Group 2; 9%) and TZP-NS (Group 5; 8%). We next looked at the minimum inhibitory concentration (MIC) distributions for each respective BL/BLI across each of the BL/BLI groups (Fig. S1B). For AMP and TZP antimicrobial susceptibility testing (AST), there was a bifurcation of susceptible and resistant isolates with very few isolates near the respective MIC breakpoints (Fig. S1B). Conversely, for SAM and AMC AST, there were large numbers of isolates which had intermediate resistant phenotypes, which likely reflects a more diffuse spectrum of resistance for SAM and AMC vs. AMP and TZP (Fig. S1B).

From the CRO susceptible *E. coli* BSI cohort, we selected 147 isolates for whole genome sequencing across the spectrum of BL/BLI susceptibilities with the group distribution of sequenced strains relative to the total cohort shown in **Fig. 1**. We under- and over-sampled Group 1 and Group 5 respectively whereas Groups 2, 3, and 4 had similar proportions of isolates sequenced within each group respective to the total cohort (148/637; 23%) as shown in Figure 1.

**FIG S1.**
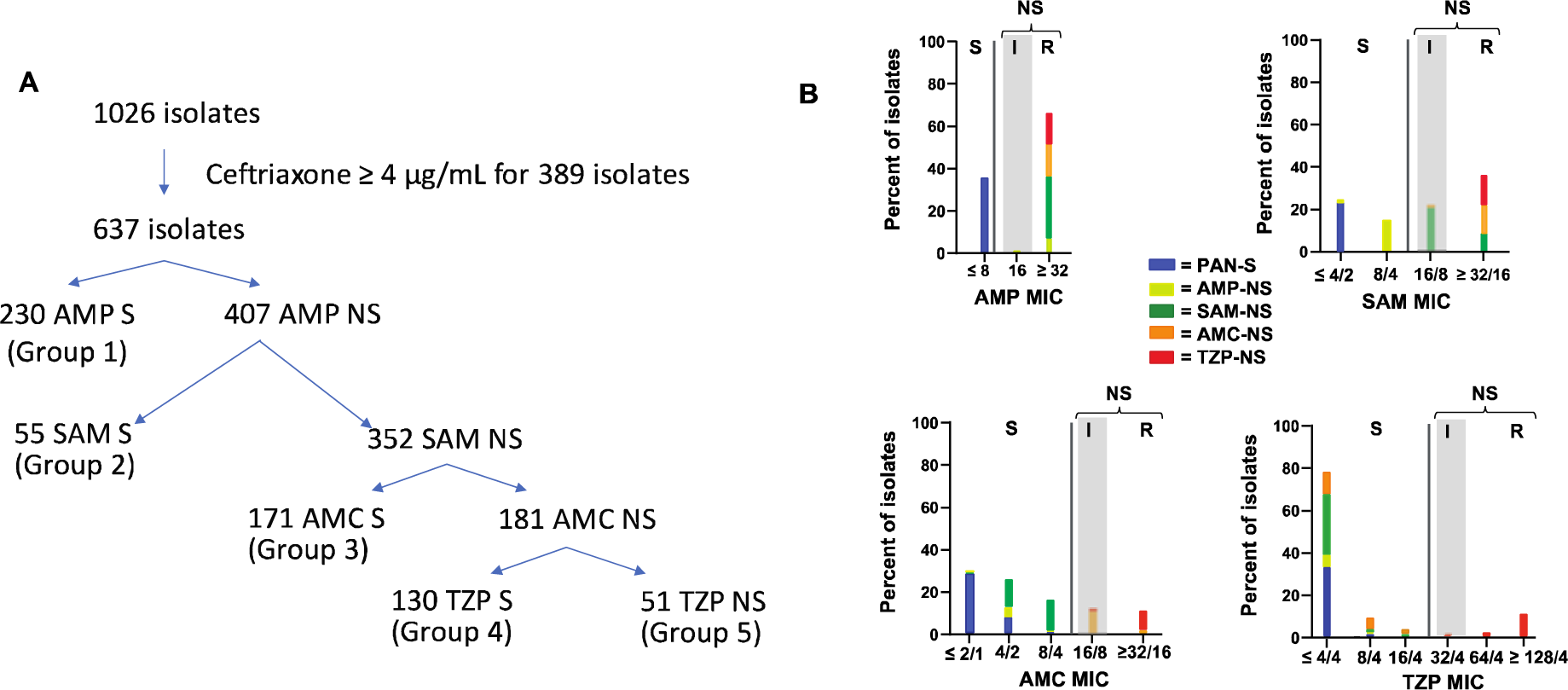
Study Population and BL/BLI MIC Distribution Across Each Non-Susceptible Group. (A) Overview of total *E. coli* bacteremia isolates detected from May 1^st^, 2015, to April 30^th^, 2020, with their respective ESRI phenotype groups. S = susceptible, NS = non-susceptible. (B) Categorical distribution of minimum inhibitory concentration (MIC; μg/mL) values across each respective BL/BLI non-susceptibility group designation for AMP, SAM, AMC, and TZP respectively. S = susceptible, I = intermediate, and R = resistant according to CLSI guidelines. Intermediate isolates are shaded gray.

### Exogenous β-lactamase gene detection contribution to BL/BLI genetic determinants

The majority of isolates which were at least AMP-NS had ≥ one exogenous β-lactamase encoding gene detected (129/134; 96%). The *bla*_TEM_ variants were the most commonly identified exogenous β-lactamase encoding genes being present in 84% (112/134) of AMP-NS or greater strains with the non-inhibitor resistant TEM (N-IRT) gene, *bla*_TEM-1B_, accounting for the vast majority (n = 110) of N-IRT detected. The *bla*_OXA-1_ gene was the second most commonly identified exogenous β-lactamase encoding gene being present in 19/134 strains (14%). Previously characterized inhibitor resistant TEM (IRT) variants were quite rare, with *bla*_TEM-31_ and *bla*_TEM-40_ each found once. Other rare β-lactamase encoding genes included *bla*_SHV-1_ (n=1), *bla*_CARB-2_ (n=2), *bla*_HER-3_ (n=1), *bla*_LAP-2_ (n=1) and two *bla*_CTX-M_ variants further described below.

For both AMP-NS and SAM-NS strains, nearly all strains carried *bla*_TEM-1_ alone with a single SAM-NS strain containing *bla*_CARB-2_ whereas *bla*_LAP_ and *bla*_CARB-2_ were present in two AMC-NS strains. Conversely, strains in the TZP-NS had the most diverse array of β-lactamase encoding genes including two strains with IRT β-lactamase variants (n = 2) and CTX-M enzymes (n = 2) respectively. One CTX-M enzyme was a derivative of CTX-M-15 (*i.e.,* CTX-M-189) with an S133G polymorphism whereas the other was a derivative of CTX-M-27 (*i.e.,* CTX-M-255) and contained a G239S polymorphism. Both of these polymorphisms have been shown in an experimental model to engender resistance to β-lactamase inhibitors but also to diminish the ability to hydrolyze cephalosporins (30). There was a statistically significant difference in exogenous β-lactamase gene content between the groups (χ^2^ *P*-value < 0.05). Specifically, AMP-NS strains were significantly less likely to contain any exogenous β-lactamase whereas TZP-NS strains were more likely to contain *bla*_OXA-1_ (Fisher’s exact test *P*-value < 0.001 for each). When only Groups 2 through 4 (*i.e.,* AMP-NS, SAM-NS, AMC-NS) were analyzed, no statistically significant differences in β-lactamase encoding gene content were observed. Thus, we conclude that the fully susceptible and fully resistant groups can be distinguished from the other groups using exogenous β-lactamase content alone, but the middle three groups cannot.

### Contribution of additional mechanisms to BL/BLI non-susceptibility

Given that exogenous β-lactamase encoding genes only separated the fully susceptible and fully resistant strains from the remainder of the cohort, we next focused on other factors previously associated with BL/BLI resistance (8). First, *E. coli* strains encode an AmpC β-lactamase which is typically transcriptionally silenced but can become active in the presence of promoter mutations (26). We identified three strains, all in AMC-NS isolates, which contained *ampC* promoter variations previously associated with AMC resistance (26). Similarly, we assessed for variation in the *bla*_TEM_ promoter region and found nine instances of strong *bla*_TEM_ promoter variants (*i.e.,* eight *Pa/Pb* and one *P5*) (24, 25). For eight of the nine cases, strains were in the TZP-NS group with the remaining strain having a borderline TZP MIC of 16 mg/L indicating that *bla*_TEM_ promoter mutations were associated with TZP non-susceptibility. Finally, we analyzed the OmpC and OmpF content of our cohort inasmuch as variation in outer membrane protein profile has been associated with BL/BLI resistance (20). Only four strains had predicted inactivating *ompC* (n = 1) or *ompF* (n = 3) mutations and the strains were present in varied groups (one in AMP-NS, one in SAM-NS, and two in TZP-NS). Thus, 16 strains (11%) had genetic changes predicted to increase *ampC* or *bla*_TEM_ expression or eliminate *ompC/ompF* production which have been previously associated with BL/BLI resistance.

### Association of *bla*_TEM_ amplification and BL/BLI resistance

Increased TEM-1 activity has previously been shown to be an integral aspect of progressive BL/BLI resistance (16, 18) with mechanisms involving increased transcription due to promoter variation and/or increase in *bla*_TEM-1_ copy number (18, 24, 25). The median copy number estimate of *bla*_TEM_ was 1.57× with a maximum of 45× observed (**Fig. 2A**). Taking a cut-off of 2.0-fold DNA coverage as evidence of amplification (31), 40/115 *bla*_TEM_ NIR containing strains (35%) had *bla*_TEM_ amplification. To understand the relationship more clearly between *bla*_TEM_ amplifications and BL/BLI resistance we focused on the 91 strains which only contained N-IRT genes as a β-lactam resistance mechanism (*i.e.,* no other exogenous β-lactamase, no *ampC*/*bla*_TEM_ promoter variation, and no *ompC/ompF* disruptions). We found statistically significant increases in *bla*_TEM_ amplification levels between SAM-NS and TZP-NS isolates (**Fig. 2B**). However, even for groups where there was a statistically significant difference, there remained overlap such that *bla*_TEM-1_ amplifications did not clearly distinguish between the various resistance profiles (**Fig. 2B**). Consistent with the inability of *bla*_TEM-1_ amplifications to distinguish between strains from AMP-NS and SAM-NS, there was no significant correlation between SAM MIC and *bla*_TEM-1_ amplification (*P*-value = 0.09, **Fig. 2C**). However, AMC and TZP MICs were significantly associated with *bla*_TEM-1_ amplification (*P*-value < 0.01, **Fig. 2D, 2E**). When *bla*_TEM_ amplification > 2× was considered as a potential β-lactam mechanism, there was a statistically significant difference in β-lactamase content between SAM-NS and AMC-NS (Fisher’s exact test *P*-value < 0.001) whereas AMP-NS and SAM-NS continued to have similar mechanisms (Fisher’s exact test *P*-value = 0.17). Thus, these data suggest that *bla*_TEM-1_ copy numbers help distinguish AMC resistant from AMC susceptible strains but do not assist with assessing SAM susceptibility.

**FIG 2.**
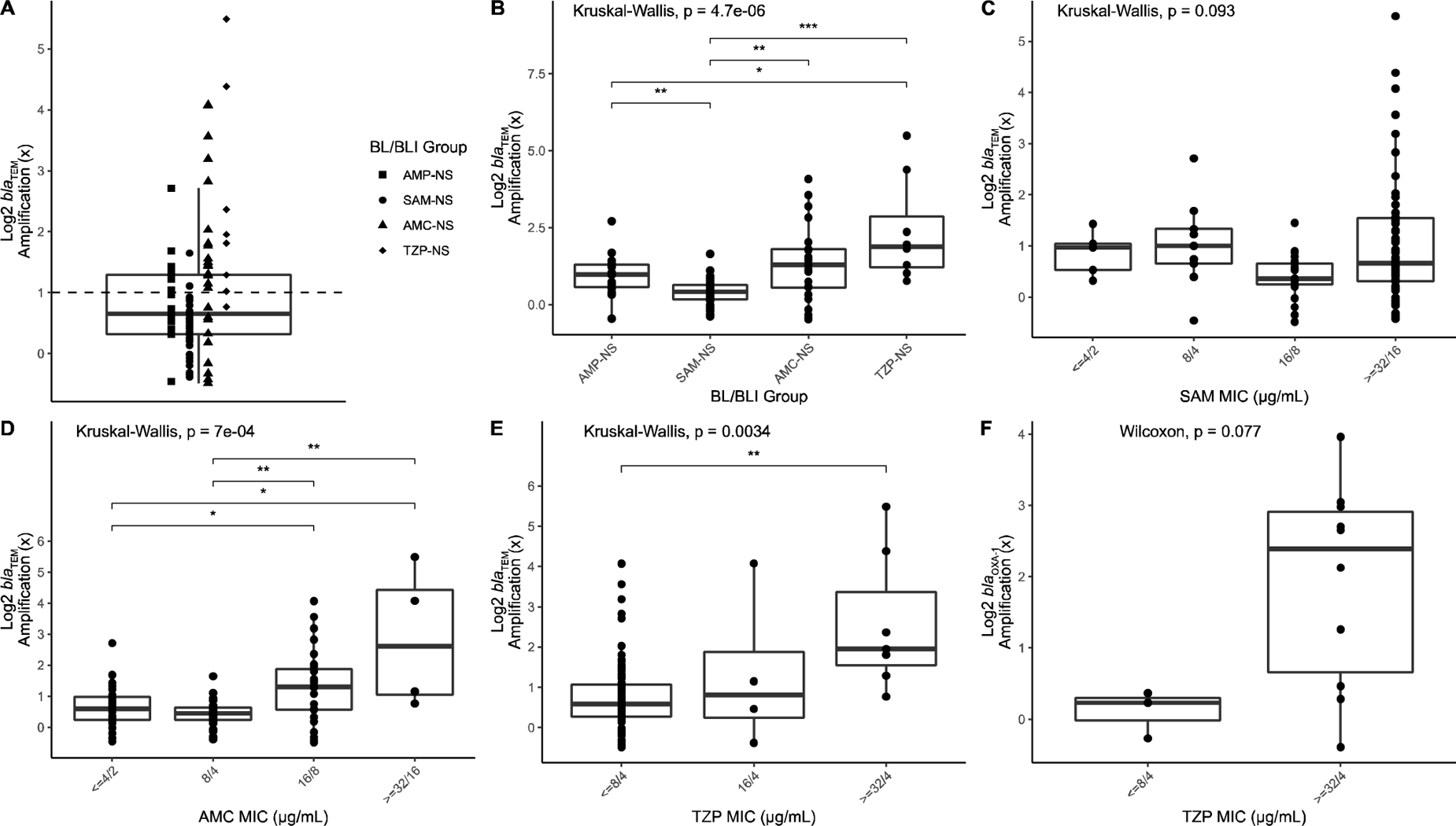
Log_2_ transformed *bla* copy number variant (CNV) estimates for BL/BLI isolates with a single exogenous β**-**lactamase gene after excluding isolates with non-exogenous BL/BLI genetic determinant factors. (A) CNV estimates for *bla*_TEM-1_ for 91 *E. coli* isolates stratified by BL/BLI group indicated by shape. Horizontal dashed line indicates cut-off for 2x CNV; (B) *bla*_TEM-1_ CNV across each of the BL/BLI groups; (C) *bla*_TEM-1_ CNV by SAM MIC; (D) *bla*_TEM-1_ CNV by AMC MIC; (E) *bla*_TEM-1_ CNV by TZP MIC; (F) *bla*_OXA-1_ CNV by TZP MIC. Global *P-*values for Kruskal-Wallis or Wilcoxon Rank-sum Tests reported above each respective panel. For global *P-*value < 0.05, pairwise Wilcoxon Rank-sum test adjusted *P*-values are reported as follows: * < 0.05; ** <0.01; *** <0.001; **** <0.0001.

As previously noted, *bla*_OXA-1_ containing strains were only present in AMC-NS and TZP-NS isolates with the vast majority of *bla*_OXA-1_ containing strains belonging to TZP-NS (16/19). The three *bla*_OXA-1_ AMC-NS strains did not contain another BL/BLI resistance mechanism whereas six TZP-NS strains contained both *bla*_OXA-1_ and *bla*_TEM-1_, and one TZP-NS strain had a TEM-promoter variation, *bla*_OXA-1_, and *bla*_TEM-1_. Similar to *bla*_TEM-1_, an increase in *bla*_OXA-1_ CNV estimates was commonly observed in our cohort with 11/19 strains (57%) having > 2.0× normalized coverage depths. When strains containing only *bla*_OXA-1_ as a β-lactam resistant mechanism were considered, there was an increase in CNV positively correlated with TZP MIC albeit this did not reach statistical significance (**Fig. 2F**). Consistent with these data, all strains with a *bla*_OXA-1_ copy number ≥ 2× were in the fully resistant category.

### Correlations of BL/BLI genetic determinants with BL/BLI groups across population structure

A core gene alignment inferred maximum-likelihood phylogeny is presented in **Fig. 3A**. There were eight core gene inferred clusters identified through the hierBAPS algorithm (32) that are highlighted in the phylogeny (**Fig. 3A**). Each of the cluster levels were associated with previously established phylogroups (33) or more closely related sequence types (STs) with highly related STs labelled on branch tips grouping together (**Fig. 3A**). The majority of isolates in this CRO-S cohort belonged to the B2 clade (60%; 88/147) although a total of 37 distinct STs were present. Strains of the pandemic ST131 clade (33%; 49/147) or the emergent ST1193 (16%; 23/147) were the most frequently observed STs with both belonging to the B2 phylogroup. There was only one other ST with more than 10 observations, which was ST69 (7%; 11/147). Other STs comprised of five or more strains included ST73 (n = 8), ST648 (n = 6) and ST10 (n = 5). We grouped STs with less than 5 strains together as “Rare STs” and such strains accounted for 31% of the cohort (n = 45). When comparing all STs comprised of ≥ 5 isolates, there was a statistically significant relationship between ST and AMR grouping (χ^2^ simulated *P*-value < 0.001) (**Fig. 3B**). Specifically, ST648 strains were more likely to be TZP-NS, and ST1193 strains were more likely to be SAM-NS (Pairwise Fisher’s Test adj. *P*-value < 0.05). Conversely, ST131 strains were not statistically significantly associated with any particular susceptibility profile. While not statistically significant, similar trends can be seen on **Fig. 3A** where ST73/ST12 isolates (hierBAPS level 7; light blue) have 85% isolates with AMC-NS or TZP-NS phenotypes whereas ST69 have more susceptible patterns with 91% of isolates having SAM-NS or a more susceptible phenotype.

**FIG 3.**
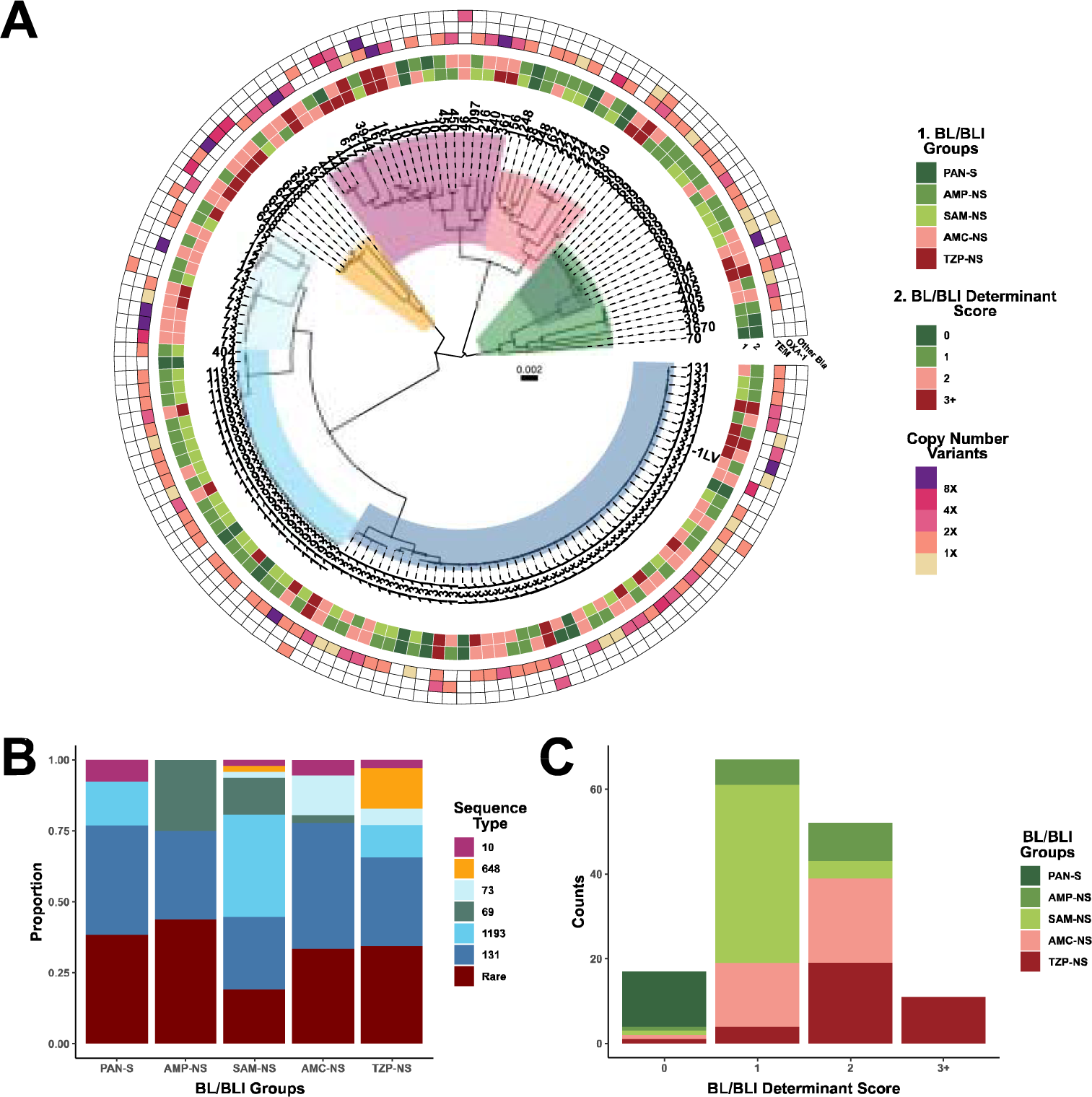
BL/BLI genetic determinant distribution across ESRI population. (A) Core gene alignment inferred maximum likelihood phylogeny of 147 CRO-S *E. coli* bloodstream Isolates. The tree is mid-point rooted with the timescale indicating mean nucleotide substitutions per site. Clades are shaded by core hierarchical population structure using the hierBAPS algorithm with each clade strongly associated with one or more phylogroup. Branch tips are labelled by respective sequence type. Inner rings 1 and 2 correspond to BL/BLI groups and BL/BLI determinant scores respectively as labelled in legend. Outer ring corresponds to *bla*_TEM_, *bla*_OXA-1_, and other *bla* gene copy number variants with copy number estimates color coded as indicated in legend. (B) Stacked bar-chart of the total proportion of STs across each of the BL/BLI Groups. (C) Stacked bar-chart of frequency counts of BL/BLI groups across each of the BL/BLI determinant scores.

We further characterized each of the BL/BLI groups by the number of BL/BLI genetic determinants, which could range from 0 to 10 (see Methods for details) (**Fig. 3C**). Each of these BL/BLI genetic determinants were based on presence/absence of genetic features that have been previously identified as contributing to BL/BLI resistance (13, 20, 24, 26, 30, 34). The number of unique combinations of BL/BLI genetic determinants across each of the BL/BLI phenotype groups are shown in Fig S2. All PAN-S isolates (n=13) had no identifiable BL/BLI determinants whereas almost all isolates (130/134; 97%) with AMP-NS or higher had at least one genetic determinant identified (Fig S2). Interestingly, all isolates in our cohort with 3+ BL/BLI determinants (n=10) were TZP-NS (**Fig. 3C**). Importantly, there was a positive monotonic relationship (ρ = 0.62, *P*-value = 2.2e-16) between BL/BLI group status and BL/BLI genetic determinant score thus underscoring how increasing numbers of BL/BLI genetic determinants impact BL/BLI non-susceptibility. Nevertheless, there was similar distribution of BL/BLI genetic determinant scores across Group 2 and Group 3 isolates (Table S1), further highlighting the difficulty in discriminating genetic differences across these isolates.

Finally, we next sought to combine genetic determinants associated with BL/BLI resistance along with our phylogenomic structure to assess the odds of being in a greater or lesser BL/BLI non-susceptibility group using ordinal logistic regression methods. **Table 1** provides an overview of BL/BLI and population level covariate relationships with the odds of being in each BL/BLI phenotypic group. There were no statistically significant associations observed between increasing BL/BLI group and ST or phylogroup; however, there was a weak association between Cluster 2, predominantly consisting of B1/C isolates that had 0.3 times (95% CI: 0.1 – 0.9) lesser odds of belonging to a higher (i.e., less susceptible) BL/BLI group compared to Cluster 1 isolates (ST131). When looking at relationships of increasing BL/BLI non-susceptibility with the composite BL/BLI genetic determinant score, for every one unit increase in BL/BLI genetic determinant, the odds of being in a higher BL/BLI group was 10.9 times (95% CI: 6.2 - 20.2) greater than a lower BL/BLI group (**Table 1**). Importantly, there were no associations between N-IRT gene carriage and BL/BLI group (OR = 1.1; 95% CI: 0.5 – 2.4) whereas *bla*_TEM_ amplification, *bla*_OXA-1_ presence, and other non-TEM *bla* amplifications all appeared to contribute to increasing odds of BL/BLI non-susceptibility. Thus, these data suggest that the presence and amplification of narrow-spectrum β-lactamases are the primary drivers of the progressive BL/BLI phenotype.

**Table 1.** Ordinal Logistic Regression to measure covariate associations with BL/BLI Groups

|  |  |  |  |
| --- | --- | --- | --- |
| **Feature** | **Mean (SD)/n (%)** | **OR<sup>a</sup>** | **95% CI<sup>b</sup>** |
| BL/BLI Genetic Determinant Score<sup>c</sup> | 1.4 (0.9) | 10.9 | 6.2 – 20.2 |
| N-IRT Presence | 110 (74.8) | 1.1 | 0.5 – 2.4 |
| *bla*<sub>TEM</sub> ≥ 2× | 40 (27.2) | 2.8 | 1.4 – 5.4 |
| *bla*<sub>OXA-1</sub> | 19 (12.9) | 33.6 | 9.0 – 125.1 |
| Other *bla* ≥ 2× | 14 (9.5) | 67.8 | 8.5 – 543.5 |
| Hierarchical Cluster<sup>d</sup> | - | - | - |
| Cluster 2 (B1/C) | 12 (8.2) | 0.3 | 0.1 – 0.9 |
| Cluster 3 (D – ST405-like) | 7 (4.8) | 0.8 | 0.2 – 3.8 |
| Cluster 4 (D – ST69-like) | 12 (8.2) | 0.3 | 0.1 – 1.0 |
| Cluster 5 (B2 – ST1193-like) | 25 (17.0) | 0.5 | 0.2 – 1.1 |
| Cluster 6 (A) | 20 (13.6) | 1.9 | 0.7 – 5.0 |
| Cluster 7 (B2 – ST73-like) | 13 (8.8) | 2.1 | 0.8 – 5.9 |
| Cluster 8 (F – ST648-like) | 8 (5.4) | 4.5 | 1.0 – 20.9 |
| <sup>a</sup>Odds ratio<br><sup>b</sup>95 percent confidence interval<br><sup>c</sup>BL/BLI genetic determinant score composite variable of ten binary coded variables (See methods)<br><sup>d</sup>Each group is compared to Cluster 1 which is B2 phylogroup ST131 isolates; parenthesis includes phylogroup and most common ST group where applicable |  |  |  |

**FIG S2.**
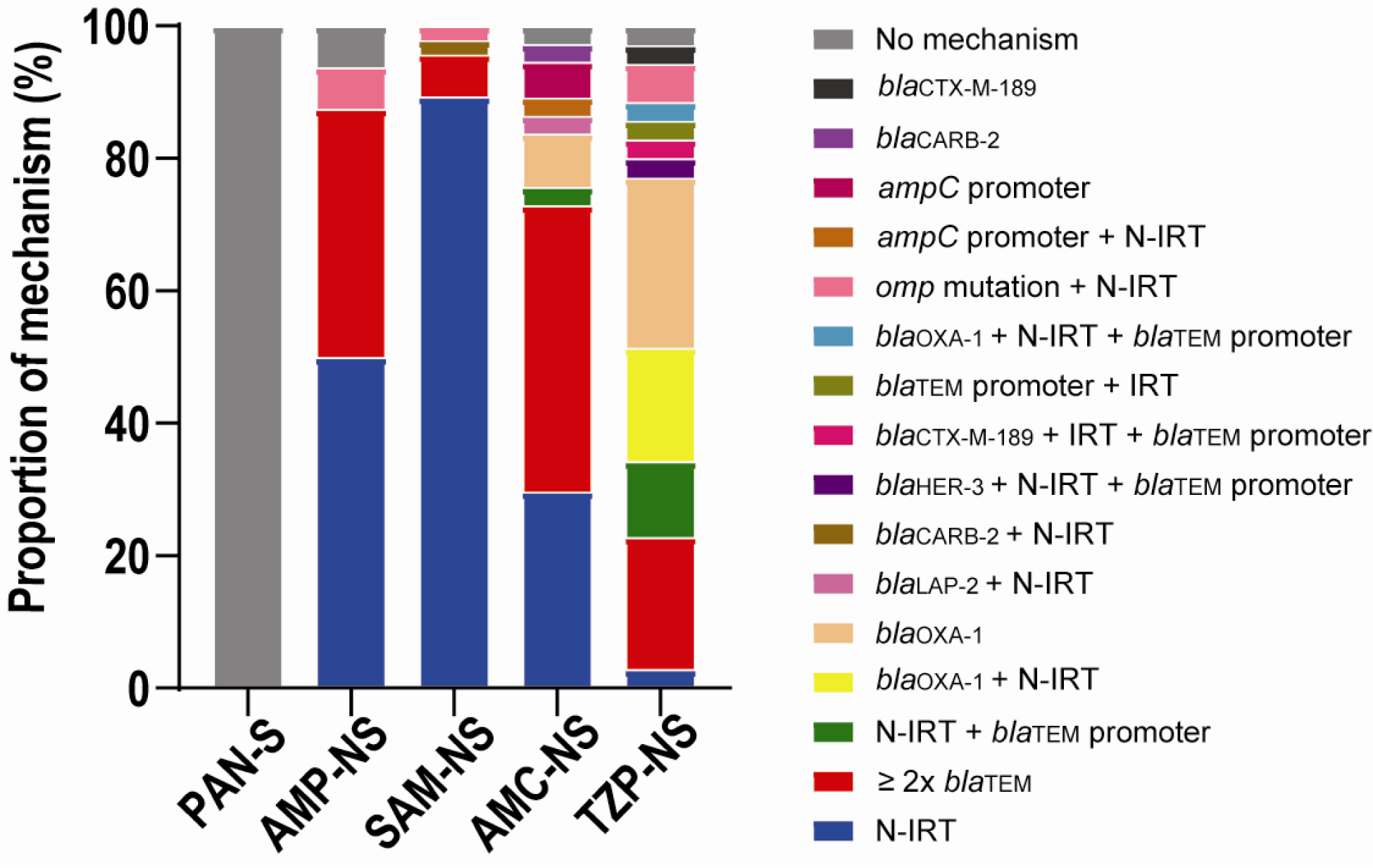
Proportion of BL/BLI genetic mechanisms detected across *E. coli* isolates by BL/BLI phenotype. Each of the respective combinations of genetic mechanisms are listed across each of the BL/BLI phenotypes and labelled accordingly in the legend. N-IRT = non-inhibitor resistant *bla*_TEM_ variant (usually *bla*_TEM-1_). IRT = inhibitor resistant *bla*_TEM_ variant, promoter = promoter variant leading to increased gene expression, ≥ 2x *bla*_TEM_ = copy number variation ≥ 2.

## Discussion

The increasing affordability and availability of WGS data has markedly increased understanding of bacterial AMR to the point where it has been postulated that such analysis might augment or even replace routine phenotypic susceptibility testing (35). For fast growing bacteria, such as Enterobacterales, WGS has shown good concordance with AMR phenotype for many “bug-drug” combinations, but in general WGS has not accurately predicated Enterobacterales susceptibility to commonly used BL/BLIs such as SAM, AMC, and TZP (5, 7, 36). Herein, we sought to determine whether a comprehensive WGS analysis, including assessment of *bla* copy number, could separate cephalosporin-susceptible *E. coli* based on BL/BLI resistance patterns. Our data indicate that an increase in copy numbers of narrow spectrum β-lactamases does occur with progressive BL/BLI resistance but accounts for only part of this process in conjunction with an accumulation of a variety of other mechanisms (13).

A key strength of our study was using WGS data to analyze a phenotypically diverse cohort of *E. coli* bloodstream isolates from across the spectrum of BL/BLI resistance in contrast to previous studies which investigated a single BL/BLI combination (8, 16, 20). Moreover, we focused on cephalosporin susceptible strains and integrated analyses of *bla* gene copy numbers to elucidate the recently proposed concept of extended-spectrum resistance to BL/BLI (ESRI) (13). This approach allowed for augmented understanding of how each β-lactam resistance mechanism contributes to the spectrum of BL/BLI resistance in clinical isolates. For example, mutations in the *ampC* promoter were associated with resistance up through AMC but not TZP whereas mutations in the *bla*_TEM_ promoter were observed only in fully resistant strains except for a single isolate with a borderline TZP MIC of 16 mg/L. Similarly, although *bla*_OXA-1_ has been associated with TZP resistance (16), we observed that strains in which *bla*_OXA-1_ was the only β-lactamase and did not have increase copy number remained TZP susceptible although resistant to AMC and SAM suggesting a necessary gene dosage and/or OXA-1 activity effect for progressive resistance. Given that *bla*_OXA-1_ amplification is clearly associated with TZP resistance (16, 17), these and other data raise concerns regarding the clinical efficacy of TZP therapy of *bla*_OXA-1_ containing strains even if they test initially susceptible given the capacity of such strains to readily increase *bla*_OXA-1_ copy number (37).

A major surprising finding of our study was that whereas there was a progressive increase in *bla*_TEM_ copy numbers moving from SAM-NS to AMC-NS to TZP-NS strains, there was conversely a small decrease in *bla*_TEM_ copy numbers between AMP-NS (Group 2) and SAM-NS (Group 3) isolates. Such data were even more surprising given that *bla*_TEM-1_ was nearly universal and the only β-lactam resistance mechanism for both groups, leading us to hypothesize that increased TEM-1 production due to gene amplification would be the major mechanism distinguishing these two groups (13). Differentiation between AMP-NS and SAM-NS *E. coli* has not been systemically studied using WGS data such as has been done for AMC-NS and TZP-NS isolates (8, 13, 15, 16, 19), and thus we are limited in our ability to benchmark our data against others. Noguchi et al. studied SAM resistant isolates from Japan using phenotypic assays and found some relationship between increased TEM-1 activity, decreased membrane permeability, and SAM resistance (20). Consistent with our data, they found only a small percentage of strains had inactivating mutations in *ompC* or *ompF*. Thus, together with other data regarding *E. coli* responses to antimicrobials, we hypothesize that *bla*_TEM-1_ harboring *E. coli* develops SAM-NS through an increase in TEM-1 activity that is not detectable via assessment of gene copy number when the organism is grown in the absence of antimicrobial pressure, along with decreased SAM entry through outer membrane changes which are not due to defined genetic mutations. As *E. coli* moves from SAM-NS to AMC-NS to TZP-NS, the resistance mechanisms then become both more diverse and genetically fixed in the population such that they become detectable using WGS based methodologies. Thus, barring identification of new genetic mechanisms that can separate AMP-NS and SAM-NS isolates, it seems unlikely that WGS methods using bacteria grown in the absence of antimicrobial pressure will be able to accurately separate these two groups. Consistent with data from other groups (8, 16), WGS analyses are likely to be useful in detecting AMC and TZP resistance in *E. coli*, but need to consider a large group of potential mechanisms, including single nucleotide polymorphisms (SNPs), which are additive in nature and thus much more difficult to predict relative to ESBL phenotypes primarily driven by single genes. There is growing interest to create ‘mechanism agnostic’ machine learning models that can classify AST phenotypes based on detecting associations with SNPs and gene presence/absence after controlling for population structure (38–41).

There are several limitations to our study worth noting. First, we sought to provide a “real-world” assessment of using WGS to analyze BL/BLI resistance and thus did not recapitulate the phenotypic data nor did we assess BL/BLI phenotypes comparing different methodologies such as broth microdilution vs. agar dilution. Thus, given the difficulties in the accuracy of BL/BLI susceptibilities using automated methodologies (23), it is likely that there was some misclassification of organisms into the various groups, particularly for those whose MICs were near the susceptibility breakpoints. Second, we used the WGS data to assess for known β-lactam resistance mechanisms and thus cannot be certain that other mechanisms were not present. Even though we assessed nearly 150 strains, our sample sizes for some groups, such as group 2, were relatively small which could have limited our power to detect statistically significant differences in various mechanisms. Finally, we only sequenced strains from a single center meaning we do not know the generalizability of our findings although our data are in accord with other data from geographically distant locales (13-15, 18, 21).

In summary, we present herein a comprehensive WGS analysis of a large cohort of cephalosporin-susceptible *E. coli* strains from across the susceptibility spectrum of commonly used BL/BLIs. Our data both support and add to the complexity of the progressive ESRI spectrum proposed by Rodriguez-Villodres et al (13). Our data suggest that the addition of β-lactam copy number and promoter variation assessment does assist with delineation of AMC-NS and TZP-NS but not SAM-NS suggesting that the initial movement of *E. coli* along the ESRI spectrum may not be detectable using methods that query AMR databases.

## Materials and Methods

### Bacterial strains and AST characterization

*Escherichia coli* bloodstream isolates are collected weekly through an IRB approved protocol and stored at −80C in 40% glycerol stocks. Initial antimicrobial susceptibility testing (AST) is performed on a Vitek®2 (bioMérieux) automated platform through the University of Texas MD Anderson Cancer Center clinical microbiology laboratory. Additional AST was performed on selected candidate isolates to determine cohort inclusion using gradient ETest strips (Liofilchem) for SAM, AMC, and TZP respectively on MHB plates. Isolates with ‘intermediate’ susceptibilities to respective BL/BLI were grouped with isolates with interpretations of ‘resistant’ within the same BL/BLI group and thus defined as non-susceptible to that particular BL/BLI combination. Further AST information is included in Table S2.

### Whole genome sequencing of bacterial isolates

Isolates with no indication of contamination and that fit into one of five groups based on AST confirmation were sub-cultured in LB and incubated for 3 hours at 37°C with 225 rpm agitation. A culture cell pellet was used to perform genomic DNA extraction using the QIAGEN DNeasy Blood & Tissue Kit following manufacturer’s instructions. The quality of gDNA was assessed through a measure of gDNA concentration with the Qubit 4 fluorometer (Invitrogen) and high molecular weight with the 4200 Tapestation (Agilent) system using the Genomic DNA ScreenTape assay. The respective QC gDNA was submitted to the MDACC Advanced Technology Genomics Core (ATGC) for library preparation and short-read, paired-end (150 × 2) whole genome sequencing using the Illumina NovaSeq6000 instrument. Samples were multiplexed, barcoded, and sequenced to achieve coverage depths > 100×.

### Short-read sequencing data QC and copy number quantification

Short-read data was QC’d and processed through a bespoke pipeline (Shropshire W, spades_pipeline, GitHub: https://github.com/wshropshire/SPAdes_pipeline). Briefly, paired-end fastq reads have Illumina adaptors and low-quality bases trimmed using Trimmomatic-v0.39 with a seed-match (16 bp) that allows up to 2 mismatches, a sliding window of 4 bp, minimum average quality = 15, and minimum length of 36 bp. Trimmed fastq reads were then quality assessed using fastqc-v0.11.9. Sequencing QC information as well as BioSample IDs are provided in Table S2.

Trimmed reads were inputted into the COpy Number Variant QuantifICation Tool (CONVICT; w Shropshire; GitHub: https://github.com/wshropshire/convict). A core-gene control file, which is created through the panaroo pan_reference.fa file, is used to normalize coverage depths. CONVICT uses a coverage-depth normalization approach whereby the coverage-depth of target genes are divided by the coverage-depth of control genes to give an estimation of ploidy-agnostic copy number. In the first step of the CONVICT algorithm, reads are aligned with a target set of antimicrobial resistance genes determined by kmerresistance (42, 43) with the resfinder database (v4.0) (44, 45). Aligned reads are then sorted with Samtools into a corresponding bam file. Coverage measurements are determined by pileup.sh from the BBMAP suite of tools in bins of 100 bases along the length of a target gene. Bins are removed or maintained in an iterative process by comparing coverage-depth of a given bin to the mean coverage-depth of all bins for the respective gene until a specific coefficient of variation (CV = 0.175) is met. Coverage-depth for control genes is calculated in the same way as for the target genes. Copy numbers are then estimated by dividing the coverage-depth of target genes by the coverage-depth of control genes. Complete convict estimated copy number results are provided in Table S2.

### Short-read assembly, database query, and phylogenetics

Trimmed reads are then used as input for short-read genome assemblies using SPAdes-v.3.15.5 using the –isolate parameter (46). Short-read assemblies were used as input to AMRfinder plus (v3.11.11) using database version 2023-04-17.1 to confirm AMR variant alleles (47) detected with the short-read based convict tool. We used known strong *bla*_TEM_ promoter variants (24) and *ampC* promoter variants (26) that have been experimentally characterized as database input into a variant annotator tool (Selvalakshmi27, VariantAnnotator, https://github.com/Selvalakshmi27/VariantAnnotator). Each variant was confirmed using blast searches as well as visually inspected on SnapGene-v5.0.8. Genome assemblies were first annotated using Prokka-v1.14.6 (48) Annotated .gff files and were used to produce core gene alignment file using panaroo-v1.2.10 (49). We used the MAFT-v7.505 aligner (50), with the panaroo parameters strict mode and a core gene threshold of greater than 99% present within the population (n=147). The core gene alignment file then was used to generate a maximum-likelihood (ML) phylogeny using IQtree2-2.2.0-beta. Parameterization included 1000 replicates for SH approximate likelihood ratio test, 1000 replicates for ultrafast bootstrap (51), and model optimization using ModelFinder (52). The optimized ML model was an unrestricted model with optimized base frequencies using a maximum-likelihood with a FreeRate of heterogeneity chosen according to BIC. Core gene alignment file was used as input to investigate population structure using an implementation of the hierarchical population clustering tool, RhierBAPS (32) with a max depth of hierarchical search =2 and max populations = 30. The ggtree-v3.3.1 R package was used for tree visualization and mapping metadata to each individual tree branch-tip (53).

### Statistics

Genetic determinants were coded into ten binary variables to create a cumulative BL/BLI genetic determinant score (Table S3). These variables included N-IRT presence, TEM gene amplification (≥2×), IRT gene, strong *bla*_TEM_ promoter variant, *ampC* promoter variant, *bla*_OXA-1_, *bla*_CTX-M_, Other *bla* gene, Other *bla* gene amplification (≥2×), and predicted Omp mutations based on previous research (13, 20, 24, 26, 30, 34). A Spearman’s rho coefficient was calculated to test for relationship between BL/BLI groups and BL/BLI genetic determinant score. Global Wilcoxon Rank Sum or Kruskal Wallis Tests were performed to detect significant differences in log transformed copy numbers between groups. For global comparisons with significant *P*-values, pairwise comparisons using Wilcoxon rank sum exact test were performed with a false discovery rate *P*-value adjustment method. Ordinal logistic regression was performed using BL/BLI group as an ordinal outcome and various genetic features as independent covariates with the MASS R package (v.7.3-54) and the ‘polr’ function with a proportional odds logistic regression model. The brant test was performed to determine if parallel regression assumption held for each model using the ‘brant’ R package (v0.3-0). Statistical analysis was performed using R-v4.0.4.

### Data Availability

Whole genome sequencing short-read data has been submitted to BioProject PRJNA924946 and PRJNA836696. R scripts for analyses can be made available through request of corresponding author.

## Acknowledgements

The MDACC clinical microbiology lab does a fantastic job identifying and transferring these organisms of interest to us and as always is much appreciated. Core grant CA016672(ATGC) and NIH 1S10OD024977-01 grant provide funding for the Advanced Technology Genomics Core (ATGC) sequencing facility at MD Anderson Cancer Center. The National Institute of Allergy and Infectious Diseases (NIAID) T32 AI141349 Training Program in Antimicrobial Resistance supports the work of W.C.S. The NIAID R21AI151536 and P01AI152999 NIAID grants support work for this project for S.A.S. The authors acknowledge the support of the High-Performance Computing for research facility at the University of Texas MD Anderson Cancer Center for providing computational resources that have contributed to the research results reported in this paper.

